# Combination Drug Therapy Reduces Iron Accumulation and Microglia-Mediated Pathologies in Neonatal Intraventricular Hemorrhage: A Biochemical and Transcriptomic Analysis

**DOI:** 10.64898/2026.02.14.705043

**Authors:** Vanessa Castro Diaz, Michelle Sunshine, Furong Hu, Sohan Shah, Weihua Huang, Carl I Thompson, Michael S Wolin, Selvakumar Subbian, Edmund F LaGamma, Govindaiah Vinukonda

## Abstract

This study describes the distribution of non-reactive brain-resident microglia densely populated along the borders of the lateral ventricles and choroid plexus in premature rabbit pups during early forebrain development. Following intraventricular hemorrhage (IVH), microglia become activated, proliferate, and migrate deeper into parenchymal regions. During this process, activated microglia exhibit a global expansion with a disproportionally elevated proinflammatory M1 nomenclature phenotype from 25% to 50% of the total; that shift was reduced by sulforaphane (SFN; Nrf2-antioxidant response element [ARE] activator of anti-inflammatory pathways) plus deferoxamine (DFN; iron chelator) treatment. Transcriptome analysis identified over expression of pro-inflammatory calcium-binding proteins S100A8 and S100A12 (intracellular damage signals), as well as chemokines CXCL8 and CXCL10 by neurons and microglia. The combination treatment of SFN-DFN mitigated M1 infiltration, suppressed the magnitude of inflammation and reduced ferroptosis after IVH in the developing postnatal brain. Moreover, SFN-DFN treatment reversed most dysregulated genes in inflammation and iron homeostasis networks, revealing potential molecular targets for additional pharmacologic interventions after IVH. We propose that reducing the toxic microcellular environment will attenuate both the injurious inflammatory responses and improve recovery of the trajectory toward normal brain development. Additionally, suppression of proinflammatory molecules and iron toxicity should promote better survival as well as salutary effects of “*living stem cell therapy*” as we have previously shown.

## 1. INTRODUCTION

Intraventricular hemorrhage (IVH) is a common problem of premature newborns and is associated with white matter injury (WMI), long-term neuromotor delay and cognitive disabilities (Volpe, 2015). Hemorrhagic extravasation of red blood cells (RBC) hemolyze and locally release toxic byproducts including free hemoglobin (heme and globin), free iron, and iron-induced free radicals contributing to an inflammatory cytokine influx and additional collateral injury (Bulbake et al., 2019). Despite advances in perinatal care over the past 30 years, a recent meta-analyses showed that the overall prevalence of IVH in extremely premature neonates (≤ 28 weeks gestational age) is unchanged at approximately 34%.(Lai et al., 2022; Nagy, Koi, et al., 2025; Nagy, Obeidat, et al., 2025). However, at the core of this phenomenon is the fact that in the earliest gestational ages, the absence of clearly effective preventative measures for IVH that directly address the structural vulnerabilities of the maturing brain, remains enigmatic (Ballabh, 2010). Moreover, no definitive therapeutic intervention exists for IVH, and clinical management is primarily supportive care. Since prevention has had limited success, we reasoned that addressing mechanisms of injury initiated at early phases of the temporal progression after IVH would likely mitigate the extent of subsequent or derivative inflammatory processes enhancing the potential for recovery of normal brain development. We speculate that preferential conditions should extend survival of “*living cellular therapy*” to better harness the evolutionary leverage of stem cells that enhance recovery from brain injury. (Vinukonda & La Gamma, 2022)

Mechanistically, IVH originates within the subependymal germinal matrix (GM), a highly vascularized and cellular brain region prominent between 22 and 32 weeks of gestation in humans (Del Bigio, 2011). The GM is a critical neurogenic niche that produces migrating neuro-glial precursor cells to form all layers of the adult brain. Hemorrhagic disruption of the GM compromises the ependymal lining allowing extension into the lateral ventricles, where severe bleeding encompasses both the sub-ventricular (SVZ) and ventricular zones (VZ) of the brain parenchyma (Finkel et al., 2023; Purohit et al., 2021). The presence of extravasated blood within these areas initiates a temporally predictable cascade of pathophysiologic events: reduced nutrient delivery due to ruptured blood vessels, extravasated red blood cells (RBCs) causing a secondary mass compression effect, then hemolysis and breakdown of hemoglobin into heme, globin and free iron. Further, the RBC lysis resulting in free hemoglobin and iron catalyzes the formation of highly toxic free radicals triggering collateral release of cytotoxic inflammatory byproducts from activated microglia that serve a dual role as essential modulators of injury and recovery. The injury progression with excessive free radical production results in a secondary late phase extension of the injury involving inflammation and apoptotic cell death. Derivative of these events is a reduction in normal growth factors that ultimately, impedes the recovery to normal brain development. (Bozza & Jeney, 2020; Dawes, 2022; Gram et al., 2014; Shao et al., 2022)

After injury resident non-reactive, microglia undergo morphological transformation into an activated phenotype, characterized by hypertrophic or amoeboid morphology with reduced process complexity. The classically activated M1 nomenclature phenotype secretes pro-inflammatory cytokines and reactive oxygen species (ROS), while the alternatively activated M2 phenotype, facilitates phagocytosis, anti-inflammatory, and tissue repair processes (Gomes-Leal, 2012). Furthermore, microglia participate in release of damage associated molecular pattern molecules (DAMPs) due to IVH pathology, which further accelerate inflammation.

The pivotal contribution of microglia to IVH pathology is not surprising as they constitute up to 10% of the total cell population in the rabbit brain and up to 16.6% in the human brain, depending on the anatomical region (Lawson et al., 1992). Moreover, in the developing human brain, resident microglia arrive from erythromyeloid precursors in the yolk sac early in the first trimester and are low in density (Andjelkovic et al., 1998; Choi, 1981; Fujimoto et al., 1989; Gao et al., 2023). By 24 weeks gestation, large numbers of microglia with short process can be identified in the germinal matrix and subventricular zone (Hutchins et al., 1990). Subsequently, microglia mature into a terminally differentiated ramified morphology and further accumulate in density in all anatomical areas until after birth. (Kostovic & Judas, 2002; Lawson et al., 1990; Perry et al., 1985; Schmid et al., 2009). Once morphology and migration are established, microglial populations are maintained via a self-renewal process throughout life with a median turnover rate of 28% annually (∼ 0.08% per day) and an estimated lifespan of 4.2 years in the human cortex (Reu et al., 2017). We hypothesized that by attenuating early-stage microglia activation after IVH injury, we can reduce the overall magnitude of the prolonged inflammatory state and improve recovery.

In this report, to gain insights into the complexity of these processes and perhaps those relevant but not yet recognized, we took advantage of RNAseq technology to elucidate changes in gene expression and associated signaling pathways in the developing brain after IVH. We attempted to redirect the pathophysiology of injury by removing free iron with deferoxamine (DFN) treatment as DFN might also reduce ferroptosis (iron-dependent apoptotic cell death). In addition, we endeavored to augment neuroprotection and to reduce cell death after IVH by mitigating inflammation with sulforaphane treatment (SFN; a naturally occurring plant Nrf2-antioxidant response element [ARE] activator that inhibits NFκB signaling). We reasoned that a genome-wide RNAseq approach would unmask iron-specific toxicity signaling in microglial gene signatures and perhaps, conditional signaling among other networks underpinning IVH pathology. We found that after IVH, SFN-DFN treatment reduced the activation of M1 microglial phenotype plus suppressed inflammation and cell death. We also delineated the transcriptomic changes associated with iron accumulation with or without DFN treatment and identified changes in iron pathway gene expression.

## 2. METHODS

### 2.1 Glycerol induced interventricular hemorrhage in premature rabbit pups

Timed pregnant New Zealand White rabbits (Oryctolagus cuniculus) were obtained from Charles River Laboratories Inc. (Wilmington, MA, USA). Premature pups were delivered via cesarean section at day 29 (E29) of gestation (term = 32 days). Newborn premature pups were maintained and fed according to protocols outlined in our previous publications (Chua et al., 2009; Georgiadis et al., 2008; Vinukonda et al., 2010; Vinukonda et al., 2019).

Intraventricular hemorrhage (IVH) was induced by administering intra-peritoneal glycerol at 4–6 hours of age, as previously described (*Georgiadis et al., 2008*). The presence of IVH was confirmed by cranial ultrasound at 24 hours of age. The severity of IVH was graded based on the echogenic volume within the ventricles, measured in three dimensions (length, width, depth) using both coronal and sagittal views. Hemorrhage was categorized into three grades: (1) no IVH: no detectable hemorrhage, (2) moderate IVH: clot volume of 100–250 mm³, with hemorrhage into the lateral ventricles and mild ventricular enlargement (distinct separation of the lateral ventricles), and (3) severe IVH: clot volume of 251–350 mm³, with marked ventricular enlargement (fusion of lateral ventricles into a single chamber) and/or parenchymal involvement.

### 2.2 Administration of pharmacological agents deferoxamine (DFN) and sulforaphane (SFN)

After ultrasound grading, pups were assigned to one of four experimental groups: (1) controls without IVH, (2) glycerol-IVH pups treated with saline, (3) glycerol-IVH pups treated with DFN, and (4) glycerol-IVH pups treated with SFN and DFN. We included 5–6 premature pups per each experimental groups for all endpoint studies. We injected pups affected with IVH with deferoxamine (DFN; Hospira, INC. IL: 60045, Cat # NDC0409-2336-01) once daily at 50 mg/kg (subcutaneously) beginning on the day of birth for 3 days (3 doses; prior to sacrifice on day 3 postnatal age). A second group of IVH-pups received 5, once daily doses and were sacrificed on day 7. When using combined treatments, pups were also administered sulforaphane (SFN; EMD Millipore # 574215) dose, 25 mg/kg intramuscular (IM) at days 1 and 3 for day 3 endpoints, and days 1, 3 and 5 for day 7 endpoint studies. Both drug schedules are based on previous publications (Cui et al., 2015; Innamorato et al., 2009; Klomparens & Ding, 2019; Sripetchwandee et al., 2014).

### 2.3 Tissue collection and processing

All experimental procedures, sample collections, and laboratory assays were approved by the Institutional Animal Care and Use Committee (IACUC) at New York Medical College, Valhalla, NY. Forebrain parenchymal tissue, cerebrospinal fluid (CSF), and plasma samples were collected from all four experimental groups (control, IVH-saline, IVH+DFN, and IVH+SFN/DFN) at postnatal days 3 and 7. Each group included 5–6 pups per time point. CSF was collected via the anterior fontanelle, snap-frozen on dry ice, and stored at −80°C for ELISA analysis. The subventricular zone (SVZ), including germinal matrix (GM), corpus callosum (CC), and corona radiata (CR), were manually dissected from 2–3 mm thick coronal brain sections (anterior to posterior midbrain) using a brain matrix slicer (Cat # BSRAS001-1, Zivic Instruments, Pittsburgh, USA). Samples were immediately snap-frozen in liquid nitrogen and stored at −80°C for RNA extraction and ELISA or real-time PCR analysis. Forebrain tissue designated for immunohistochemistry (IHC) was fixed in 4% paraformaldehyde, embedded in optimal cutting temperature (OCT) compound, and coronally sectioned from 1.0 mm anterior to 1.0 mm posterior to bregma.

### 2.4 Histological evaluation of hematoma and Iron accumulation

Hematoxylin and eosin (H&E) staining and iron staining were performed on fixed coronal forebrain sections using previously established protocols (Vinukonda et al., 2019). We assessed the distribution of hematoma and iron deposition in the SVZ in all experimental groups. Every 20th section was evaluated, resulting in approximately 10 serial sections per rabbit pup.

### 2.5 Quantification of free Hemoglobin and Iron deposition

SVZ tissue samples were lysed according to manufacturer methods. Protein concentration in the tissue lysates and CSF was quantified using the Pierce BCA Protein Assay Kit (Cat #23225, Thermo Scientific). Free hemoglobin was measured using the Abcam hemoglobin assay kit (Cat # ab272533, Abcam, USA), and total iron content was determined using a colorimetric iron assay kit (Cat # ab83366, Abcam, USA). Quantification was performed according to each kit’s manufacturer instructions.

### 2.6 Immunofluorescence staining (IHC)

Immunohistochemistry was performed as previously described (Finkel et al., 2023; Purohit et al., 2021; Vinukonda et al., 2010). Fixed forebrain coronal sections were rehydrated in 0.01 M PBS, incubated overnight at 4°C with primary antibodies, followed by incubation with secondary antibodies at room temperature for 1 hour. Sections were mounted using SlowFade™ Light Anti-fade Reagent (Molecular Probes, Invitrogen, CA, USA) and visualized under a fluorescence microscope. Primary antibodies used were: (1) Iba-1 (Cat # ab5076, Abcam) to detect total microglia, (2) Anti-Ki-67 (Cat # M724029-2, DAKO, Denmark) to assess cell proliferation, and (3) MHC class II/CD40 (Cat # 136-00042-025, Bay Biotech) to identify M1 microglia. All sections were counterstained with DAPI. TUNEL staining was performed to assess microglial apoptosis in combination with Iba-1 labeling.

### 2.7 Cell density and quantification procedures

Fluorescent images of the SVZ (including GM, CC, CR) were captured at 40× magnification after staining with Iba-1 (total microglia), MHCII/CD40 (M1 microglia), and TUNEL (apoptotic cells). Two to three alternate sections per pup were analyzed, with 5–6 pups per group. TUNEL-positive cells were quantified following the manufacturer’s protocol (Apoptosis Kit Cat # 17-141, EMD Millipore/Sigma, St. Louis, MO, USA). Two investigators, blinded to group assignments, independently performed cell counts using ImageJ software with a grid overlay and their results were averaged. Data are presented as mean ± standard error of the mean (SEM).

### 2.8 RNA isolation, RNA sequencing and TaqMan assays for gene expression

Total RNA was extracted from SVZ dissected tissue using the Qiagen Total RNA Isolation Kit (Cat # 74104, Qiagen, USA). RNA quantity and integrity were assessed using a NanoDrop® Spectrophotometer ND-2000C (Thermo Fisher, Waltham, MA, USA). RNA sequencing was performed with ERCC (External RNA Control Consortium) spike-in controls added immediately after tissue collection for normalization and detection limit assessment. Libraries were prepared using the Illumina TruSeq Stranded mRNA Library Prep Kit and sequenced on the Illumina NextSeq-550 platform (paired-end, 2 × 75 bp) at the Genomics Core Laboratory at New York Medical College. The raw RNA-seq data are published and available in the GEO database (Accession # GSE185438 and GSE198497).

Gene expression and pathway analyses were conducted using the Partek Genomics Suite (ver.6.5) and Ingenuity Pathway Analysis system (Qiagen, CA, USA) respectively - after alignment and analysis of RNA-seq data. A false discovery rate (FDR) of 5% was applied to normalized data to identify differentially expressed genes (DEGs), which were used in IPA pipeline for network and pathway analysis as reported previously (PMID:22645653; 34732811). Differential gene expression and pathway/network analyses were performed in collaboration with the Genomics Core Facility at Rutgers New Jersey Medical School.

### 2.9 Statistical analysis

The rabbit pups were assigned randomly to the experimental groups. Scatter plots and bar graphs represent mean ± SD or mean ± SEM as described in the figure legends. TaqMan assays were performed in duplicate for each sample. Pairwise differences between groups at each postnatal age were assessed using t-test and Mann-Whitney test. Group comparisons at individual postnatal ages were analyzed using one-way ANOVA using GraphPad Prism 6 (GraphPad Software, CA, USA). *Post hoc* comparisons were conducted using Tukey’s multiple comparison test. A p-value of <0.05 was considered statistically significant.

## 3. RESULTS

### 3.1 Simultaneous administration of Sulforaphane (SFN) and Deferoxamine (DFN) results in reduced hematoma and iron accumulation in the sub-ventricular region of developing brain after IVH in premature rabbit pups

To assess whether IVH leads to hematoma and iron deposition in the germinal matrix (GM), subventricular zone (SVZ), and periventricular zone (PVZ), we conducted histological analyses using our well established glycerol-induced premature rabbit pup model of IVH (Chua et al., 2009; Georgiadis et al., 2008; Vinukonda et al., 2010; Vinukonda et al., 2019). The coronal forebrain tissue blocks were dissected along the anterior-posterior axis to include both lateral ventricles and the SVZ, following our published protocols and established methods for neurotoxicological evaluation in rabbits (Pardo et al., 2020; Purohit et al., 2021). As shown in Figure 1A-F, upper panel, (H&E), iron (blue) and hematoma (brown) deposits were observed in the GM and SVZ regions of IVH pups over postnatal day 3 (Figure 1A vs B) and day 7 (Figure 1D vs E). Whereas no such deposition was seen in healthy controls at either postnatal days. Of note the iron deposition increased by postnatal day 7 in IVH pups, suggesting progressive accumulation. Importantly, after SFN-DFN treatment this accumulation was suppressed at both postnatal ages (Figure 1B, E vs C, F).

**FIGURE 1:**
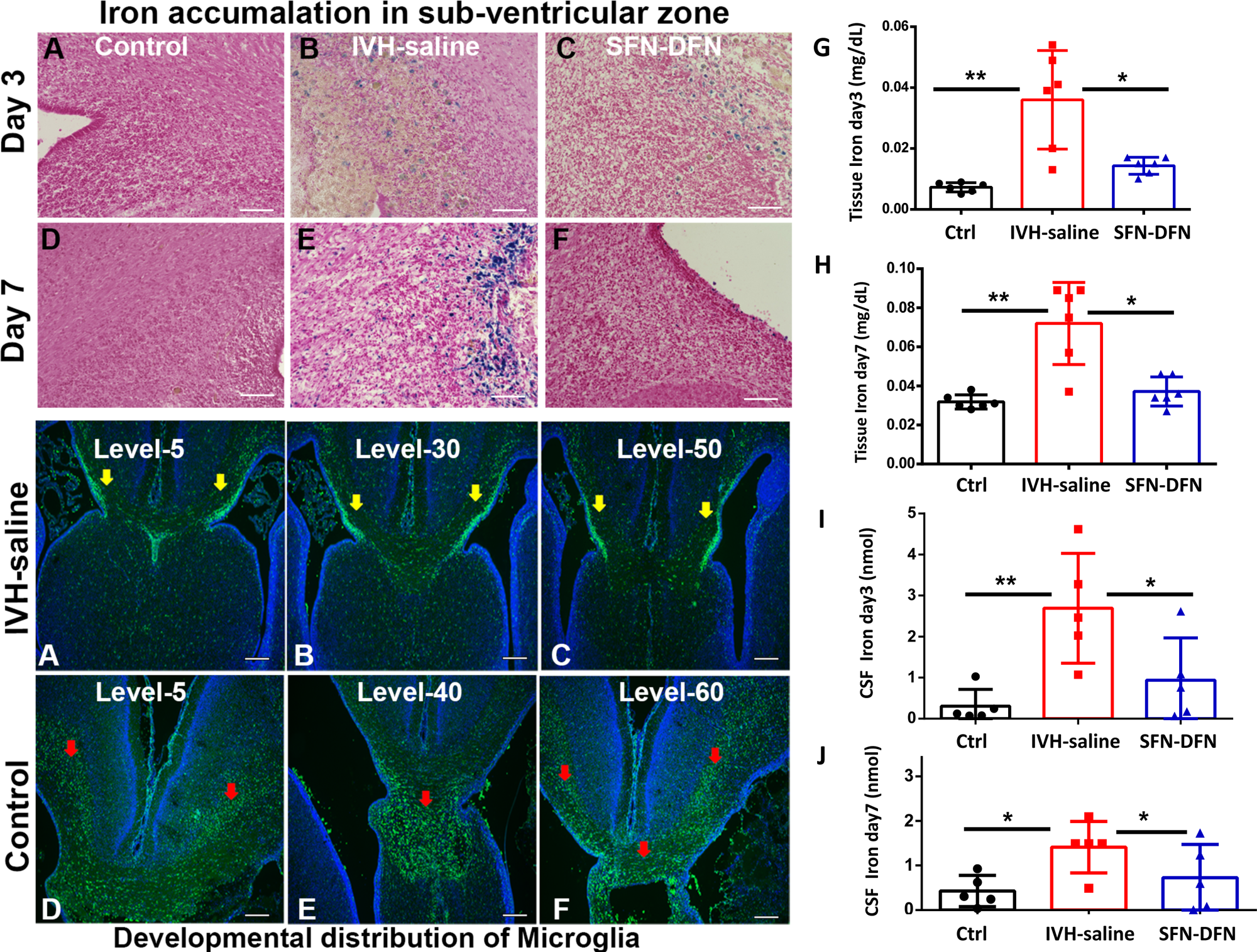
Hematoma/iron accumulation and developmental distribution of microglia in subventricular zone (SVZ) in healthy controls and after intraventricular hemorrhage (IVH) in premature rabbit pups. **A-C, Upper Panel)** Representative hematoxylin and eosin staining (H&E) on coronal section images showing hematoma and iron accumulation in the sub-ventricular regions of germinal matrix (GM) in healthy control and pups with IVH at postnatal day 3. The images indicate hematoma (brown) and iron (blue) accumulation after IVH (A vs B). No hematoma or iron deposition was seen in healthy control at postnatal day 3 (A). The IVH resulted in hematoma/iron accumulation that was reduced after SFN-DFN treatment at postnatal day 3 (C). **D-F)** Representative hematoxylin and eosin staining (H&E) on coronal section images evaluating hematoma and iron accumulation in the sub-ventricular regions of the germinal matrix (GM) in healthy control and pups with IVH at postnatal day 7. The images indicate hematoma (brown) and iron (blue) accumulation after IVH (D vs E). No hematoma or iron deposition was seen in healthy controls at postnatal day 3 (D). The IVH resulted in hematoma/iron accumulation that was reduced after SFN-DFN treatment at postnatal day 7 (F). **A-C, Lower Panel)** Representative immunofluorescence-stained images on coronal serial sections using IBA-1^+^ antibody staining (total microglia marker) in the sub-ventricular zone and ventricular borders in healthy control pups at postnatal day 3. Developmentally dense distribution of microglial immunoreactivity was observed in the healthy controls. **D-F, Lower Panel)** Representative immunofluorescence-stained images on coronal serial sections using IBA-1^+^ antibody staining (total microglia marker) in the SVZ and ventricular borders in premature pups with and without IVH injury at postnatal day 3. Postnatal developmental distribution of resident microglia primarily appeared as densely packed cells around the ventricle borders in healthy controls (A-C). Whereas after IVH resident microglia infiltrated and dispersed away from the ventricle borders into deeper parenchymal areas (D-F) at early injury by postnatal day 3. The coronal serial sections of forebrain were taken from anterior to posterior regions and used for immunostaining (H&E and IHC). Images show multiple serial sections spaced 20-30 sections apart and matched anatomical regions in control and IVH pups. The images were taken using a Keyence microscope (Keyence Corporation of America, Illinois, USA). The scale bar for all images 20 µm. **G-J)** Sulforaphane (SFN) and Deferoxamine (DFN) treatment diminished IVH-induced iron accumulation in tissue lysates and CSF during postnatal days 3 and 7 in premature rabbit pups. **G-H)** Scatter plot bar graph shows SVZ dissected tissue lysates for free iron accumulation and clearance after SFN-DFN treatment in premature rabbit pups at postnatal day 3 (G) and for postnatal day 7 (H). The data presented for tissue lysate as mg/dL with S.D. Sample size of 5-6 in each group for both day 3 and 7, *P < 0.05; **P < 0.01 were considered as significant. One way ANOVA for day 3, F(2, 15) =13.64; p=0.001) and for day 7, F(2, 15) =12.66; p=0.001). **I-J)** Sulforaphane (SFN) and Deferoxamine (DFN) treatment diminished IVH-induced iron accumulation in CSF during postnatal days 3 and 7 in premature rabbit pups. CSF collected from each experimental pup was used to assess iron levels. Total iron was assessed according to method described in the kit protocol. The data presented as nmols with S.D. Sample size of 5-6 in each group for both day3 and 7, *P > 0.05, **P > 0.01 were considered as significant.

Further, there was a significant increase in free iron levels in the dissected SVZ tissue lysate after IVH compared with healthy control pups at day 3 (Figure 1G, p<0.01, n=6 in each group) and day 7 (Figure 1H, p<0.01, n=5-6 in each group); this deposition was significantly reduced after SFN-DFN treatment at both postnatal days 3 (Figure 1G, p<0.05, n=6 in each group) and day 7 (Figure 1H, p<0.05, n=5-6 in each group).

Similar changes were observed for iron quantification in CSF collected from the three experimental groups at postnatal day 3 and 7 (Figure 1 I-J, p<0.05, n=5 in each group for both days 3 and 7). This data indicated that after IVH a significant accumulation of free iron occurred in the SVZ and that after SFN-DFN, this accumulation was reduced.

### 3.2 Developmental distribution and effect of IVH on resident microglia at postnatal day 3

To assess the response to IVH of brain resident macrophage-microglia after free hemoglobin (fHb) and iron accumulation, we examined the density of Iba-1^+^ resident microglia in the CNS and the rate of microglial cell division using Ki-67^+^ dual staining (Figure 1A-F lower panel, IF, serial sections). In healthy controls, as expected, dense populations of Iba-1 positive microglia were observed along the ventricular borders, septal region (between the lateral ventricles), and choroid plexus versus lower microglial density in the white matter and cortex (Figure 1A & C, lower panel). On the other hand, after IVH, the densely populated microglial multiply several fold and are found further away from the ventricular borders deeper into the brain parenchyma (Figure 1D-F, lower panel, IHC, serial sections). On day 3, Iba-1 (Figure 2 A-B) and dual staining with Iba-1 and Ki-67 antibodies identified total microglia population and those proliferating cells (Ki-67^+^) in coronal sections. Healthy control pups exhibited a typical ramified microglial morphology with multiple processes (Figure 2A; inset, 40x high magnification) where few microglia were positive for Ki-67 immune reactivity, indicating minimal proliferation (Figure 2C; blue arrow). In contrast, following IVH, the infiltrated microglia acquired an amoeboid morphology and spread into deeper parenchymal regions (Figure 2B, inset, high magnification). These infiltrated microglial cells were almost all positive for Ki-67 indicating strong mitotic activity (Figure 2D; blue arrows). These data indicate that after IVH, resident microglia are activated and become highly mitotic and appear to have migrated deeper into the parenchymal regions of the SVZ, CC and CR regions.

**FIGURE 2.**
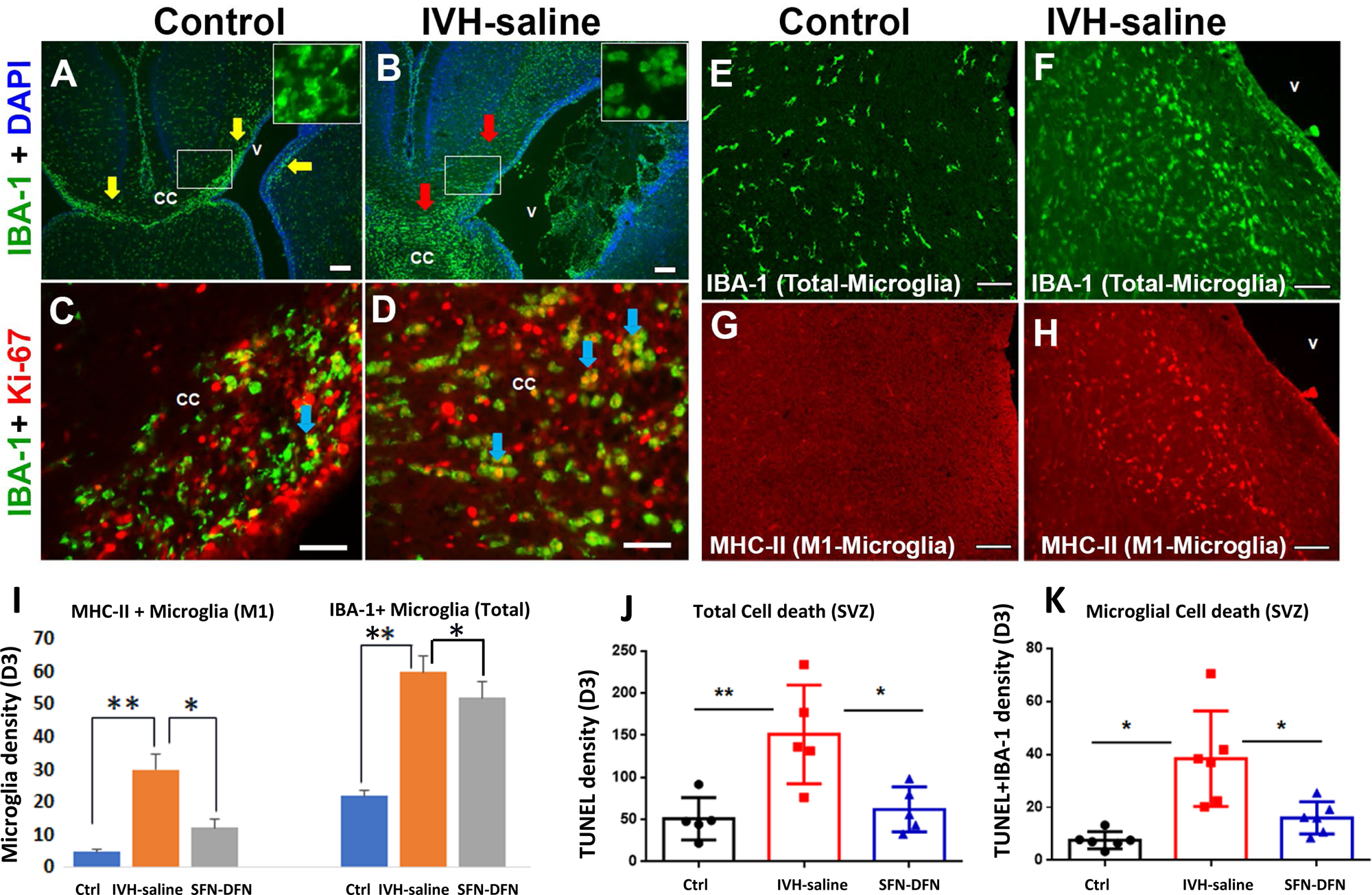
**A-L)** Developmental distribution and effect of IVH on resident microglial phenotypic change and effects of SFN-DFN treatment on IVH in premature rabbit pups. **A-D)** Representative immunofluorescence images showing microglia changes after IVH. A) Iba-1 positive total microglia form dense tracks (immune signal in green) in the ventricular borders and corpus callosum (CC) in healthy controls [ramified morphology (inset-A), less proliferative and show few co-localized cells in yellow color (C)] at day 3. B) After IVH, Iba-1^+^ cells now appear dispersed deeper into the parenchyma (SVZ and PVZ), show amoeboid morphology (inset-B), are highly proliferative shown as more co-localized dividing cells in yellow (D). Yellow and Red arrows show microglial tracks and dispersal deeper into the parenchyma (A, B). Blue arrows show proliferating microglia, co-localized, Iba-1 with Ki-67 (C, D). The low magnification images compare Iba-1^+^ immune signals for control and IVH (A-B) and at high magnification show Iba-1^+^ combined with Ki-67^+^ (proliferation marker; C-D) in premature pups at postnatal day 3. **E-H)** Representative immunofluorescence staining on coronal sections showing total and activated M1 microglia at day 3. The coronal sections were stained with Iba-1 (total) and MHC II (M1 specific proinflammatory) antibodies. The Iba-1 positive (non-reactive, ramified) were few and no MHC II positive immune cells were seen in healthy controls (E vs G). After IVH, a large number of microglia were positive for MHC II (red) indicating M1 microglia (F vs H). Scale bar 20µm, Ventricle (v), corpus callosum (CC). **I)** Cell density quantification after SFN-DFN treatment suppressed M1 infiltration induced after IVH at postnatal day 3. Bar graphs show significant increase in M1 (MHC-II positive, Figure2 I, left panel) and total microglia (Iba-1^+^, Figure 2 I, right panel) cell density compared with healthy controls. This increased density was reduced after SFN-DFN treatment at postnatal day 3. The cell count was performed on coronal section images taken in 20X magnification in both left and right hemispheres (counts include GM, CC and CR sub-regions). The data presented mean and SD in each group. Sample size of 5-6 pups in each experimental group, **P < 0.01, *P < 0.05 considered as significant. **J-K**) Scatter plot with bar graph shows cell density quantification for total cell death (TUNEL positive cells) and specific microglia cell death (Iba-1^+^ + TUNEL double positive cells). The data indicated a significant increase of total and microglia specific cell death in IVH compared to controls at both postnatal ages day 3 and 7 (J, K). Whereas after SFN-DFN treatment the total cell and microglial specific cell death were suppressed at both postnatal days 3 and 7 (J, K). The cell count was performed on 20X image. The data presented mean absolute cell count with SD. Sample size of 5 pups in each group, *P < 0.05, **P < 0.05 considered as significant.

### 3.3 SFN-DFN treatment suppressed proinflammatory M1 microglia infiltration and cell death induced by IVH in the SVZ of the developing brain in premature pups

Immunofluorescent staining of total microglia (Iba-1^+^) and a M1 specific marker (MHC-II^+^) indicated that ∼50% of activated microglia were M1 phenotype in IVH compared to about 25% in no IVH healthy controls at baseline (Figure 2I, 2E, G vs F, H). Cell density quantification (Fig 2 I) showed some effect of SFN-DFN on the rise in total microglia (Iba-1^+^) after IVH (Fig 2 I); however, SFN-DFN specifically reduced the number of M1-phenotype proinflammatory microglia vs IVH alone (MHC-II^+^; *p<0.01, n=5 each group). TUNEL staining indicated a significant increase in total cell death in IVH compared with control which was decreased after SFN-DFN treatment compared with IVH alone at postnatal day 3 (Figure 2J; p<0.01, n=5 in each group for both comparisons). Similarly, dual staining TUNEL positive cells + Iba-1^+^ microglia showed significantly reduced microglia specific cell death after IVH when treated with SFN-DFN (Figure 2K; p<0.05, n=5 in each group for both comparisons).

### 3.4. Assessment of IVH induced transcriptome changes and effects of DFN treatment

To uncover global gene expression changes in IVH and to identify iron-regulated pathways affected by DFN treatment, we analyzed the transcriptomic profiles of SVZ dissected forebrain tissues. Our RNA-seq raw data are publicly available in the GEO database (Accession # GSE185438 and GSE198497).

#### 3.4.1 Principal component analysis

We first determined the distribution of variance in the RNAseq data of control versus IVH samples using principal component analysis (PCA). The PCA revealed distinct clustering of data between control and IVH groups (Figure 3A), consistent with a clear segregation of gene expression profiles between IVH and control groups. Similarly, after administration of DFN treatment, another distinct clustering pattern was observed among IVH (no treatment, control) and DFN treatment groups, though data from one sample each of the DFN and control groups remained as outliers (Figure 3B).

**FIGURE 3.**
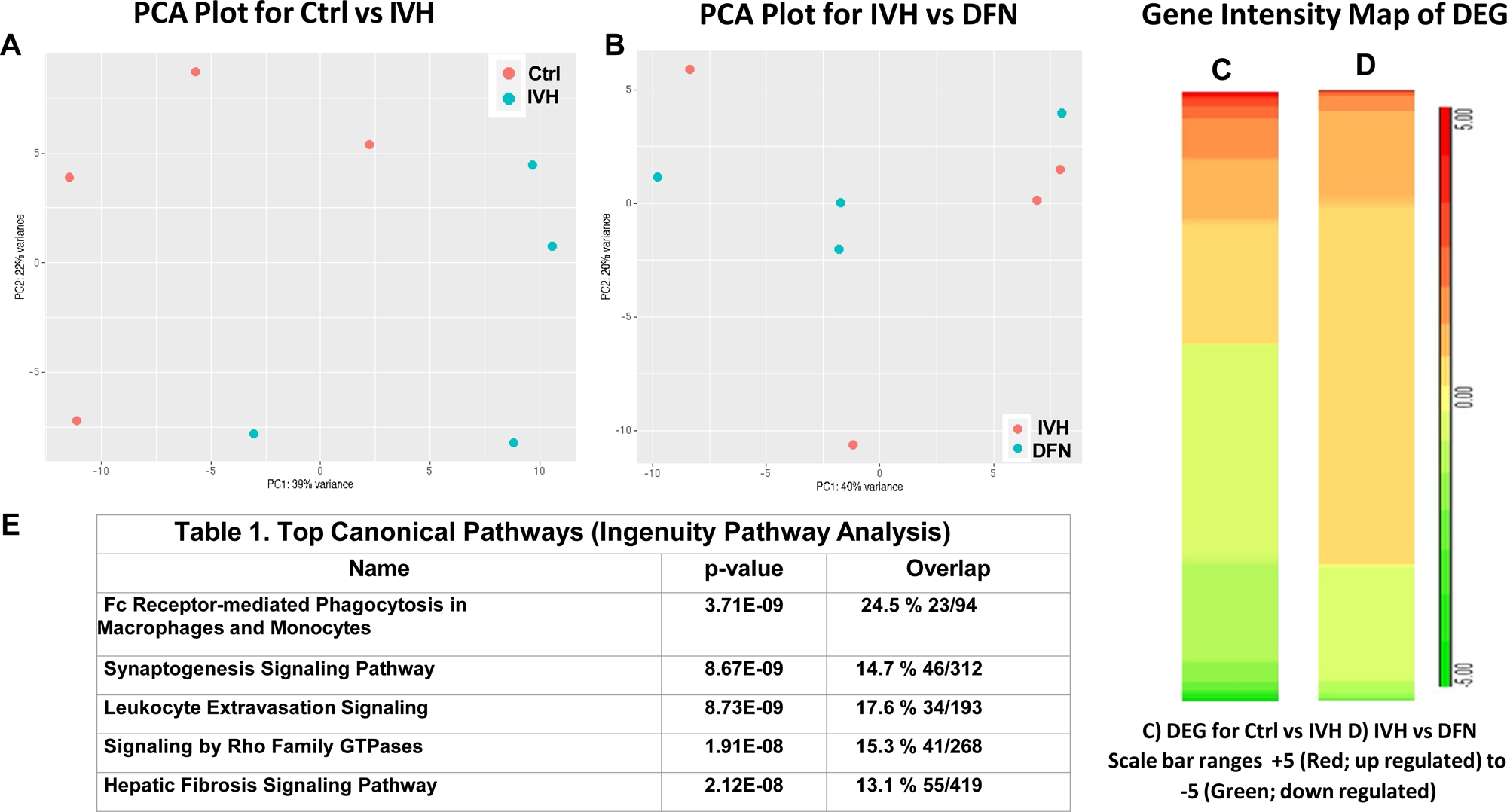
Genome-wide transcriptome changes in IVH and the effect of DFN treatment on gene expression in dissected SVZ tissue at postnatal day 3 premature rabbit pups. **A-B)** Principal component analysis (PCA) of RNAseq data highlights specific segregation and clustering of different experimental groups at postnatal day 3 (Ctrl: control, IVH; DFN-treated). The RNAseq data of samples (n= 3-4 per group) were distinctly distributed between healthy controls vs IVH (A) and IVH vs DFN treated pups at day 3. The controls samples (red colored circles) formed a separate cluster independent of IVH (blue colored circles) based on their global gene expression profiles (A). We do not see any overlay of red and blue circles gene expression profiles. The PCA plot between IVH vs DFN treated samples also visualized separate clusters (B). Note that in (B), there was one outlier of each IVH and DFN data sets, which were excluded from further analysis (see main text). The label on X and Y axes show PC1 and PC2 variances respectively. Total RNA isolated from SVZ dissected tissue was used for RNAseq. PCA analysis was performed as per standard pipeline processing of RNAseq data at the Genomics Core Laboratory at New York Medical College. **C-D)** Intensity map of differentially expressed genes (DEG) in IVH and after DFN treatment at postnatal day 3. Heat map of control vs IVH (C) indicated a total of 1,373 DEGs in IVH at postnatal day 3 and 440 DEGs after DFN (D) treatment. Color intensity map in IVH visualized brighter (red intensity for upregulation and green for downregulation) with multiple blocks compared with controls (C); both red and green color intensity was reduced after DFN treatment (D). Data were derived from n= 3-4 pups per group. Note: The intensity of color in C and D is proportional to the degree of gene expression. Scale bar ranges from +5 (Red; upregulated) to −5 (Green; downregulated). Gene expression data presented in C and D corresponds to Log2 mean fold change. **E)** Top Canonical pathways identified by Ingenuity pathway analysis. Table showing the most significant canonical pathways across the entire data sets. The highest percentage of gene expression change related to phagocytosis function in macrophage and monocytes and leukocyte extravasation signaling in SVZ dissected RNA sequencing data at postnatal day 3. The significance indicates the probability of association of molecules in the study with the canonical pathway data sets in IPA, which utilizes Fishers Exact test for P-value calculations.

#### 3.4.2 Analysis of differentially expressed gene Intensity maps and top Canonical Pathways

The profile of significantly differentially expressed genes (DEGs) was visualized using heat intensity maps. IVH pups exhibited 1373 DEGs compared with no IVH controls, while DFN treatment modulated 440 of these genes, compared to no treatment controls (Figures 3C and 3D). The gene intensity map and the number DEGs indicate that expression of about 30% of IVH dysregulated genes were impacted by DFN treatment. Further, the Ingenuity Pathway Analysis (IPA) of DEGs revealed several significantly dysregulated canonical pathways after IVH compared to controls, particularly in the innate immune signaling pathways such as Fcγ-receptor-mediated phagocytosis and leukocyte extravasation cascades (Figure 3E, table). Moreover, 23 genes out of 94 in the phagocytosis pathway are related to microglia and macrophages (Supplementary Table 1). These findings highlight the extraordinary inflammatory nature of IVH and the role of microglial phagocytic activity.

#### 3.4.3 Differentially regulated cellular functions in IVH, and the effect of DFN treatment

Based on our initial observation of significant changes of gene expression in IVH pups compared with healthy control pups, we sought to determine the cellular biological functions that were affected by the DEGs. As shown in Figure 4A, we observed upregulation of immune responses, particularly in innate cells, such as phagocytosis, inflammation and recruitment of activated leukocytes in IVH samples compared to controls (Figure 4A). Consistently, cellular functions associated with morbidity or mortality as well as tissue-level death were also upregulated in IVH. Thus, the cellular functions dysregulated in IVH are consistent with the pathological sequelae seen in these pups. Next, we investigated the cellular biological functions effected by DFN treatment after IVH, compared to no treatment controls. In the DFN-treated pups, cellular functions associated with activation and recruitment of immune cells were dampened (Figure 4B) and, consistently, cell survival function pathways were upregulated. Furthermore, neuronal development pathways were also downregulated, reinforcing the importance of immune regulation during brain development. Taken together, many of the inflammatory response-related functions that were upregulated and contributed to elevated tissue injury in IVH were downregulated after DFN treatment.

**FIGURE 4.**
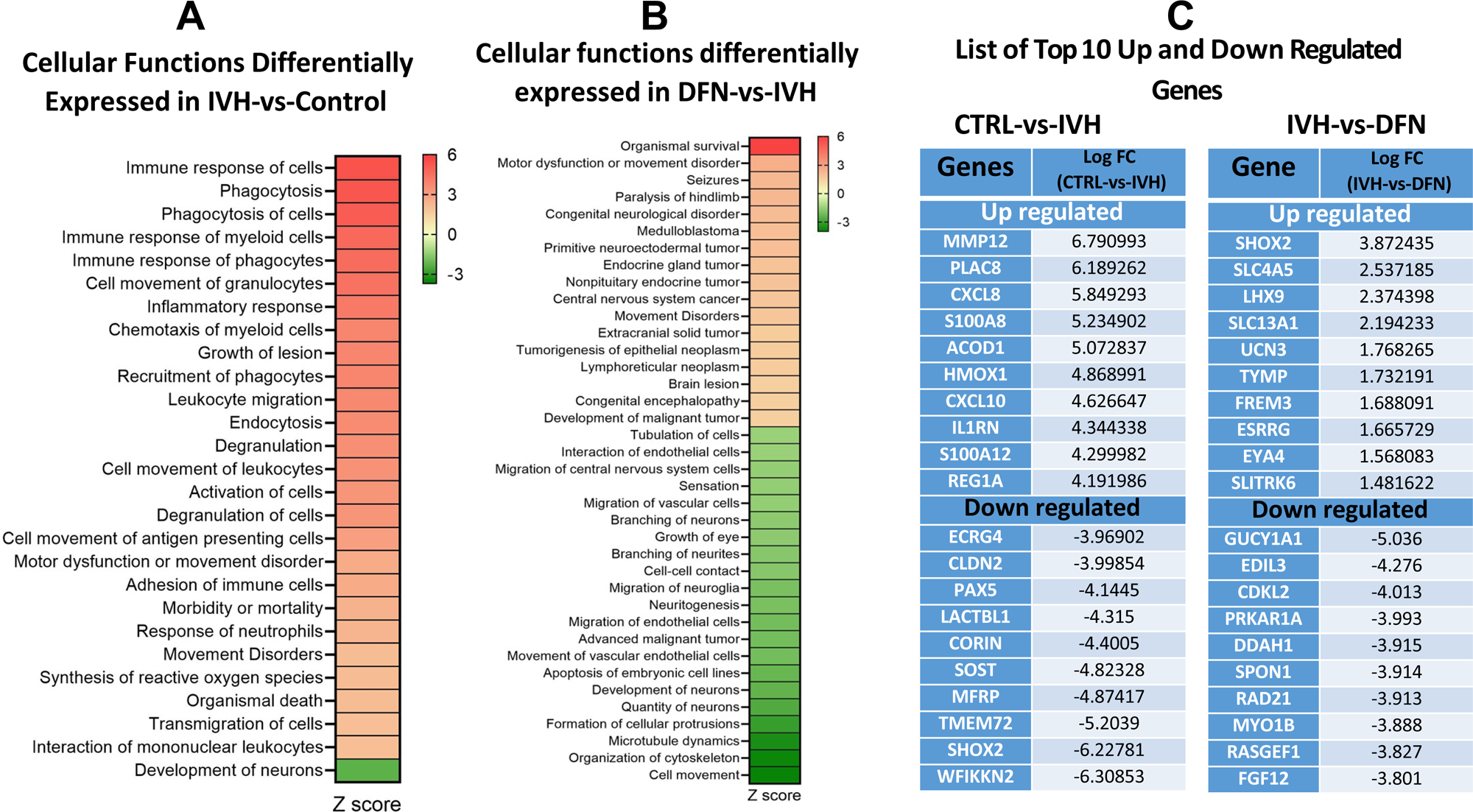
Differentially regulated cellular functions and list of highly up regulated/down regulated genes in IVH and the effect of DFN treatment in premature rabbit pups. **A-B)** Heat map of differentially regulated cellular biological functions in IVH compared to controls (A) and in DFN treated versus no treatment groups (B). IPA analysis of DEGs shows strong up regulation of cellular response of immunity, phagocytosis and inflammatory related genes in IVH, compared to controls (A). However after DFN treatment many of the IVH- upregulated responses were downregulated (B). Color intensity indicates the magnitude of significance and the color scale is an indicator of significance based on z-scores. Red color indicates upregulation and green color indicates downregulation of cellular function. Data derived from n= 3-4 pups per group. Scale bar ranges from +6 (Red; upregulated) to −3 (Green; downregulated). **C)** List of top 10 up- and downregulated DEGs in IVH and effect of DFN treatment. The column on the left show’s gene symbols of up- and downregulated DEGs and their log2 fold change in IVH compared to control group (4-fold or higher is listed). The right column depicts up- and downregulated (Log2 fold change) DEGs in IVH vs DFN treated groups at postnatal day 3. The abbreviations of gene symbols shown in the table are expanded in Supplemental Table 2.

#### 3.4.4 Highly dysregulated genes in IVH and during DFN or SFN-DFN treatments

To further extract biological insights on specific genes that are involved with the pathophysiology during IVH, and after DFN treatment, we determined the top 10 most up- and downregulated DEGs and compared them between experimental groups. The list of the top ten upregulated genes in IVH, compared to controls (all >4-fold) included: CXCL8, CXCL10, S100A8, S100A12 HMOX1, MMP12, PLAC8, ACOD1, IL1RN and REG1A. The down regulated genes in this group (ranging from 4- to 7-fold decrease) included: ECRG4, CLDN2, PAX5, LACTBL1, CORIN, SOST, MFRP, TMEM72, SHOX2 and WFIKKN2 (Figure 4C). The top 10 most significantly upregulated DEGs in DFN treated pups, include transcriptional regulators (SHOX2, LHX9, ESRRG, EYA4), neuronal signaling (UCN3, TYMP, FREM3) and ionic balance (SLC4A5, SLC13A1), while the downregulated DEGs are associated with endothelial function (EDIL3, SPON1, MYO1B), nitric oxide-cGMP/GTP signaling (GUCY1A1, DDAH1, PRKAR1A, RASGEF1, CDKL2), and regulation of organ development (RAD21, FGF12). In sum, the list of top up- and down-regulated genes in IVH (compared to no-IVH) and DFN treatment (compared to no-treatment) are consistent with their role in altering corresponding cellular functions shown in Figure 4A-B.

To confirm and validate some of the findings of RNAseq data on top 10 up- and downregulated genes in Figure 4C, we used TaqMan gene expression assays (Figure 6C-F) and we confirmed elevated expression of CXCL8, CXCL12 and S100 genes (S100A8 & S100A12) in IVH compared to controls. These over expressed chemokines are known to regulate their biological effects via CXCR1-3 receptor signaling (Zhou et al., 2023). S100A8 & A12 proteins also regulate TLR signaling mechanisms (Donato et al., 2013). Both types of ligands share MAP kinase (MAPK) signaling components to elicit distinct pathological effects on microglial activation to cause inflammation, apoptosis and other effector functions (Foell et al., 2007; Tao et al., 2022; Zhou et al., 2023).

#### 3.4.5 Identification of differently expressed Canonical Pathways and Microglial network genes and effects of DFN Treatment

To gain additional insights into the canonical pathways impacted by the DEGs of IVH and/or DFN treatment, we performed IPA Canonical pathway analysis. Our analysis revealed upregulated immune-related processes in IVH, including TREM1 signaling, MIF signaling, and reactive oxygen and nitric oxide production (Figure 5A). In contrast, neuronal pathways like cAMP, CREB, synaptogenesis, and thrombin signaling were down regulated. Importantly, DFN treatment down regulated many of the inflammatory response associated pathways including NFkB signaling, neuroinflammation signaling, Wnt/Ca pathway, P2YR signaling and CREB signaling in neurons (Figure 5B).

**FIGURE 5.**
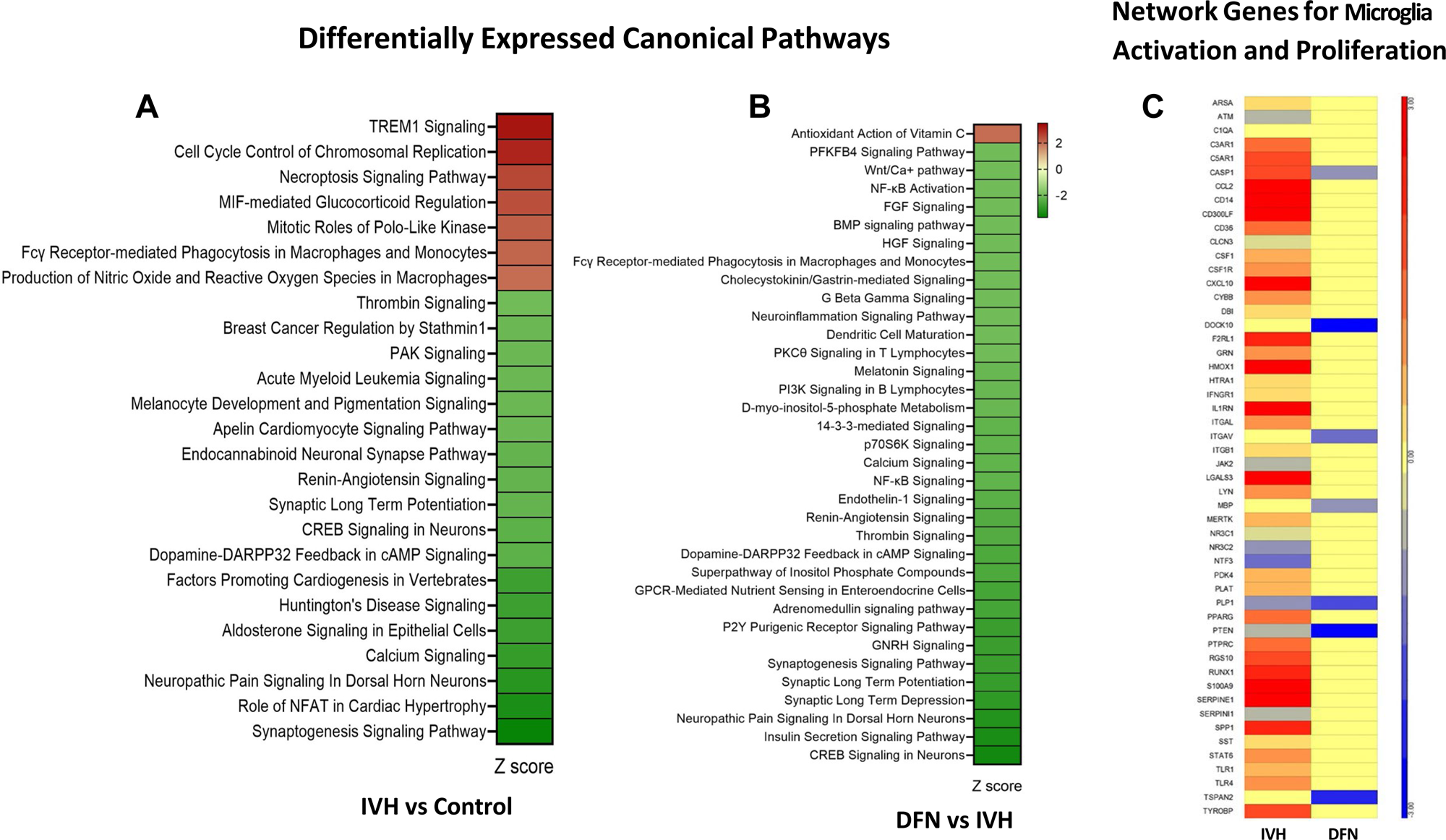
Differentially regulated canonical pathways and microglial activation and proliferation network gene expression in IVH and the effect of DFN treatment. **A-B)** Heat maps of differentially regulated canonical pathways in control vs IVH (A) or DFN treated versus no treatment (B) at postnatal day 3. Color intensity indicates the magnitude of significance and the color scale is an indicator of significance based on z-scores. Red color indicates upregulation and green color indicates downregulation of cellular function. Data derived from n= 3-4 pups per group. Scale bar ranges from +2 (Red; upregulated) to −2 (Green; downregulated). **C)** Heat map representing differentially expressed genes (DEGs) involved in microglia activation and proliferation in IVH and treated with DFN at day 3. Left column for IVH and right column for DFN treatment. Row indicates each gene as labeled and color-coded comparison between columns for each gene in row. Data derived from n= 3-4 pups per group. The intensity of color is proportional to the degree of gene expression. Scale bar ranges from +5 (Red; upregulated) to −5 (Blue; downregulated). Gene expression data corresponds to Log2 mean fold change. **Note:** Data indicated large number of microglia activation and proliferation network genes over expressed in IVH but most of these genes were downregulated after DFN treatment. The abbreviations of gene symbols shown in the table are expanded in Supplemental Table 2.

Gene network analysis highlighted that most of the affected genes in IVH are involved in microglia activation and participate by inducing inflammation, free radical production and cell death (Figure 5C). Consistently, microglial network genes involved in inflammation and cell death included: CASP1, DOCK10, ITGAV, TSPAN2, and PTEN that were upregulated in IVH, while the expression of each of these genes was down-regulated after DFN administration (Figure 5C).

#### 3.4.6 Assessment of ferroptosis network genes expression in IVH and effect of DFN treatment

Iron toxicity causes cell death via mechanisms known as ferroptosis. We identified significant upregulation of a key marker of ferroptosis (HMOX1; Figure 5C). We further explored this in the form of heat maps to identify interacting partner genes involved in ferroptosis networks after IVH and compared the effects of DFN treatment (Figure 6A, variable red color intensity). Key upregulated genes in the ferroptosis network in IVH included: HMOX1, CTSB, FTL, PRM2, LPCAT1, and CDK1. In contrast, DFN treatment dampened these gene expressions, compared to no treatment group. Importantly, expression of multiple key genes involved in iron metabolism and transport function included: ACSL4, TFRC, SLC7A11 and ABCA4 which were all downregulated in IVH and this trend was reversed after DFN treatment (Figure 6A). Further analysis of the interaction among ferroptosis-linked genes indicated that HMOX1 is the key molecule that connects several others, including p38 MAPK (cell survival/death) and ACSL4, TFRC, FTL (heme-iron regulating) pathways in IVH (Figure 6A and B). Related to this, up- regulation of CDK1 and CDKN1A would activate CTSB, a protease involved in initiating cell death and was observed in IVH pups. Importantly, expression of genes in this network were down regulated upon DFN treatment. Taken together, these findings underscore a role for DFN in modulating ferroptosis, particularly through HMOX1, and thus, possibly protecting developing brain tissue from ferroptosis after IVH. Future studies can address individual mechanisms using siRNA and knockout/ knock-in *in vitro* and *in vivo*.

**FIGURE 6.**
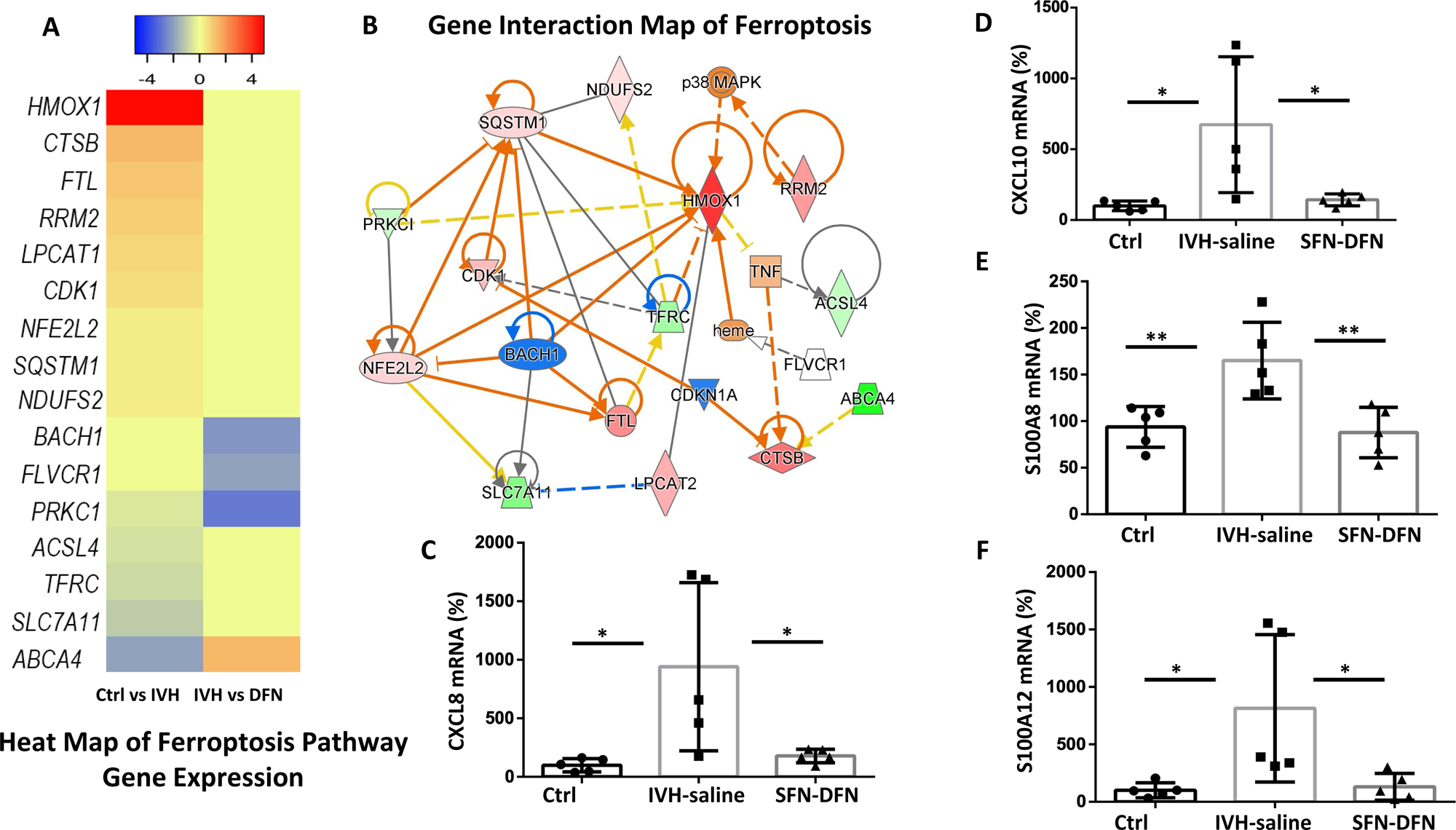
Heat map of ferroptosis network genes/pathway mechanisms and effect of DFN treatment and TaqMan assay confirmation of highly dysregulated genes in IVH and DFN treatment. **A-B)** Heat map of ferroptosis pathway gene expression in IVH and treatment (A) and ferroptosis network interaction map (B). The expression level and list of DEGs involved in ferroptosis network in IVH vs Control (left column) and effects of DFN treatment, IVH vs DFN (right column). Row indicates each gene as labeled and color-coded comparisons between columns for each gene in a row. Data derived from n= 3-4 pups per group. The intensity of color is proportional to the degree of gene expression. Scale bar ranges from +4 (Red; upregulated) to −4 (Blue; downregulated). Gene expression data corresponds to Log2 mean fold change. **Note**: Data indicated over expression of ferroptosis genes IVH and this trend was suppressed after DFN treatment. B) Image showing gene interactions among DEGs in ferroptosis networks. The color-coded symbols indicate functions of the transcript in interacting pathways. The intensity of color is proportional to the degree of gene expression. Red color indicates upregulation and green color indicates downregulation. Solid lines indicate direct interactions, and dotted lines indicate indirect interactions. The abbreviations of gene symbols shown in the table are expanded in Supplemental Table 2. **Note**: Data indicated over expression of HMOX1 is a key molecule connected to P38 MAPK (Cell survival/death), while CDK1 and CSTB protease initiates cell death in IVH. Similarly, altered heme regulated pathway molecules like ACSL4, TFRC and FTL indicated iron homeostasis imbalance in IVH that was corrected to normalcy by DFN treatment. **C-F)** SFN-DFN treatment suppressed selected inflammatory and chemokine gene expression that were induced by IVH at postnatal day 3. The bar graphs show quantitative TaqMan gene expression using total RNA isolated from the SVZ of premature rabbit pups at day 3 and effects of SFN-DFN treatment. **C-D)** The mRNA expression levels for chemokine CXCL8 (C) and CXCL10 (D) were significantly increased in IVH compared with healthy controls and this increase was significantly reduced after SFN-DFN treatment at day 3. Sample size= 5 pups per group, *P < 0.05 for all comparisons. One way ANOVA for CXCL8, F(2, 12)= 6.211; p=0.01) and for, F(2, 12)= 6.563; p=0.01). **E-F)** The mRNA expression levels of calcium binding proteins S100A8 (E) andS100A12 (F) were significantly increased in IVH compared with healthy controls and this increase was significantly reduced after SFN-DFN treatment at day 3. Sample size= 5 pups per group, *P < 0.05 (D &F); **P < 0.05 (E) for comparisons. One way ANOVA for S100A8, F(2, 12)= 9.518; p=0.003) and for S100A12, F(2, 12)= 5.691; p=0.01).

## 4. DISCUSSION

It is well recognized that the recovery from perinatal IVH is complex and multifactorial involving a temporal dependence of both early and late injury mediators contributing to the disruption of postnatal brain development. In this report we showed that a combination therapy of SFN-DFN after IVH can: i) reduce free iron in the SVZ germinal zone and circulating CSF, ii) mitigate the shift to inflammatory M1 microglia, iii) reduce apoptosis/ferroptosis and iv) mitigate the rise of a wide range of microglial related proinflammatory genes and gene networks as identified using RNAseq analysis. This proof-of-concept data suggests that early treatment with a pharmacologic intervention shortly after IVH can alter the course of injury and recovery by reducing the cascade of inflammatory responses arising from extravasation and lysis of red blood cells. It is plausible that these effects can lessen the hostile microenvironment during recovery from the incident injury and help recover processes relevant to the normal cascade of brain developmental. Moreover, we speculate that these salutary effects can extract a more favorable microenvironment effect for “living cellular therapy” to be effective when using stem cell interventions as these cells would also be less likely to die.(Vinukonda & La Gamma, 2022)

Initial histological analysis for hematoma/fHb and free iron accumulation showed that these deposits were significantly reduced after SFN–DFN combination treatment (Figure 1, upper panel, H&E). Moreover, under normal conditions, microglia densely populate the lateral ventricular borders in the subventricular zone (SVZ), and choroid plexus borders (Figure 1, bottom panel). After IVH, they increase mitosis and cell density, thus appearing to have also migrated into the parenchyma, and then ultimately, differentiate predominantly as the proinflammatory M1 phenotype; a sequence that was attenuated by SFN-DFN treatment (Figure 2E-I). Morphologically, these microglia exhibited an amoeboid shape and rounded cell bodies, consistent with a pro-inflammatory phenotype and a heightened activation state (Figure 2A-D (Colonna & Butovsky, 2017; Song & Colonna, 2018; Wang et al., 2023)). During this critical developmental window, microglial activation is known to contribute to the release of pro-inflammatory cytokines and reactive oxygen species, exacerbating injury. Importantly, apoptotic microglia cell death (IBA-1^+^ + TUNEL double positive cells) significantly increased after IVH yet, following SFN-DFN treatment, cell death was significantly reduced (Figure 2J-K). Taken together, our findings indicate that combined SFN-DFN treatment effectively reduces free iron accumulation, which was associated with fewer microglia transitioning to the M1 pro-inflammatory phenotype but to a lesser degree on the population increase of the M2 phenotype (Figure 2I, right panel).

Maintaining a balanced state between microglial activation, proliferation, and apoptosis is critical for normal postnatal brain development. Microglial proliferation and activation peaked on postnatal day 3 after IVH compared with no IVH. At that time, RNAseq data revealed significant alterations in gene expression profiles among IVH pups compared with healthy controls and in IVH compared with DFN monotherapy treated pups (at postnatal day 3). In the IVH group, 1,373 genes showed altered expression, while DFN treatment reduced that number to 440 differentially expressed genes, indicating a substantial modulatory effect. Notably, after IVH, among these changes were genes related to Fcγ-receptor-mediated phagocytosis in microglia (Figure 3).

Cell death can be influenced by factors such as lack of oxygen (e.g. due to mass effect or disruption of blood flow after IVH), glutamate excitotoxicity, or exposure to blood components like free iron. Our gene network analysis revealed that the top 10 up/ down regulated genes involved upregulated transcripts (>4-fold) for key chemokines such as CXCL8 and CXCL10, along with damage-associated molecular pattern (DAMP) molecules S100A8 and S100A12 in IVH (Figure 4 & 6). Elevation of genes involved in heme metabolism (e.g., HMOX1) and ferroptosis pathways further highlighted the complex, multifaceted nature of IVH pathology. Importantly, DFN treatment significantly downregulated the expression of these genes by postnatal day 3, suggesting early therapeutic efficacy and a potential window of opportunity as a therapeutic intervention.

Canonical pathway analysis identified activation of genes related to microglial functions, including inflammation, free radical generation, and cell death signaling. DFN treatment restored many of these pathways toward baseline expression levels (Figure 5A-B). Additionally, genes related to microglial proliferation and activation were markedly upregulated in IVH and normalized with DFN treatment (Figure 5C). Collectively, these findings provide strong transcriptomic evidence of dysregulated innate immune and phagocytic responses to IVH where DFN treatment offers partial restoration of homeostatic gene expression patterns in microglia.

Our findings also indicated ferroptosis-related gene changes in the SVZ, driven by interactions among key regulators for ferroptosis, oxidative stress and inflammatory genes such as: HMOX1, TNF, CTSB, TFRC, FTL, and components of the NRF2/p38 MAPK pathway (Figure 6). These results suggest that iron removal alone is insufficient to address the complex etiology of IVH-related injury and abnormal brain development.

Future implications of this work challenge the traditional focus on mono therapies targeting individual proinflammatory cytokines, growth factors, use of steroids, or single molecules in various pathways. Unfortunately, while instructive, former approaches yielded limited translational clinical successes (Wróblewska & Swietliński, 2005), **o**ne notable exception is the DRIFT protocol (Drainage, Irrigation, and Fibrinolytic Therapy). DRIFT demonstrated modest long-term benefits in neonates but was halted prematurely due to the risk of secondary perinatal hemorrhage (Luyt et al., 2020; Whitelaw et al., 2010). On the other hand, the significance of DRIFT findings underscored the importance of a multitiered approach directed at early interventions that concurrently address hematoma/iron clearance and attenuation of secondary inflammatory injury mechanisms arising from hemolysis-derived toxic mediators. In the current report, our results afford a new strategy using less invasive, pharmacologically driven tactical approaches (compared to DRIFT) to aid recovery from IVH.

Limitations of these experiments include the need for further validation of changes identified by RNAseq analysis using siRNA, gene knockout, over expression and single cell transcriptomic strategies to more specifically address effects on individual cell types. In addition, this proof-of-concept approach of transcriptome analysis needs independent quantification of the magnitude of RNA transcripts plus evidence that changes result in their cognate proteins. Lastly, a more expanded analysis of drug dosing, timing, and combinatorial interactions is likely to be instructive on how best to exploit a combinatorial approach to benefit brain recovery.

## 5. CONCLUSIONS

This study is the first to describe the developmental distribution of non-reactive brain-resident microglia densely populated along the borders of the lateral ventricles and choroid plexus in premature rabbit pups at early forebrain development on postnatal day3. Following IVH, these microglia become activated, proliferate, and appear to migrate into deeper parenchymal regions of the injured brain. Notably, the activated microglia also exhibit polarization into a proinflammatory M1 nomenclature phenotype but with a diminished proportion of the M2 protective phenotype - cells that are critical for tissue repair and resolution of inflammation and other developmental interactions in the developing brain. Importantly, SFN-DFN treatment mitigated M1 transformation, resulting in an increased ratio of M2 microglia that was also associated with reduced cell markers of inflammation and apoptotic cell death in the developing postnatal brain.

Furthermore, this study is the first to report in the context of IVH, dysregulation of the Fcγ receptor–mediated phagocytosis pathway, a key component of innate immune function in microglia and macrophages. Transcriptomic analysis identified over expression of S100A8 and S100A12, pro-inflammatory calcium-binding proteins that serve as intracellular damage signals, as well as CXCL8 and CXCL10, potent chemokines expressed by neurons and microglia that act as key chemoattractants during neuroinflammation. Together, these findings offer insights into microglial dysfunction, innate immune network gene pathway dysregulation following IVH, and illustrate improvements after SFN-DFN combined treatment. *We speculate* that reduced inflammation will enhance the possibility of recovering normal brain maturation (Vinukonda et al., 2019) and that reducing endogenous cell death may also increase the likelihood of stem cell survival (living cellular therapy) plus stem cell salutary effects while adapting to a changing local microenvironment (Vinukonda & La Gamma, 2022).

## Acknowledgements

We acknowledge scientific and editorial review by Bistra Nankova, PhD and secretarial support from Rita Daly in this work.

## Data Availability

The raw data generated from each experiment, which support the findings presented in this study, are available from the corresponding author and senior authors upon reasonable request, in accordance with institutional policies.

---

**Supplementary Table 1.** FcgR mediated phagocytosis in microglia and macrophages.

| **Symbol** | **Entrez Gene Name** | **Expression Ratio**<br>**(IVH-vs-no IVH)** |
| --- | --- | --- |
| HMOX1 | Heme oxygenase 1 | 4.869 |
| RAC2 | Rac family small GTPase 2 | 2.296 |
| FYB1 | FYN binding protein 1 | 1.863 |
| SYK | Spleen associated tyrosine kinase | 1.688 |
| VAV1 | Vav guanine nucleotide exchange factor 1 | 1.588 |
| ARPC1B | Actin related protein 23 complex subunit 1B | 1.443 |
| LCP2 | Lymphocyte cytosolic protein 2 | 1.394 |
| INPP5D | Inositol polyphosphate-5-phosphatase D | 1.208 |
| HCK | HCK proto-oncogene, Src family tyrosine kinase | 1.103 |
| ACTA2 | Actin alpha 2, smooth muscle | 1.028 |
| LYN | LYN proto-oncogene, Src family tyrosine kinase | 0.811 |
| WAS | WASP actin nucleation promoting factor | 0.811 |
| PLD2 | Phospholipase D2 | 0.7 |
| VAMP3 | Vesicle associated membrane protein 3 | 0.585 |
| ACTC1 | Actin alpha cardiac muscle 1 | 0.429 |
| ARPC4 | Actin related protein 23 complex subunit 4 | 0.258 |
| PIK3R1 | Phosphoinositide-3-kinase regulatory subunit 1 | -0.308 |
| PRKCI | Protein kinase C iota | -0.514 |
| PLD5 | Phospholipase D family member 5 | -0.517 |
| PTEN | Phosphatase and tensin homolog | -0.586 |
| VAV3 | Vav guanine nucleotide exchange factor 3 | -0.753 |
| PRKCQ | Protein kinase C theta | -0.91 |
| EZR | Ezrin | -1.259 |

**Supplementary Table 2.** Expansion of Gene annotation used in the study.

| **Gene Symbol** | **Gene Name** |
| --- | --- |
| MMP12 | Matrix mettalloproteinase-12 |
| PLAC8 | Placenta-specific protein-8 |
| CXCL8 | Chemokine (C-X-C motif) ligand 8 or IL-8 |
| S100A8 | S100 calcium-binding protein-A8 |
| ACDO1 | Aconitate decarboxylase-1 |
| HMOX1 | Heme Oxygenase-1 |
| CXCL10 | Chemokine (C-X-C motif) ligand-10 |
| IL1RN | Interleukin-1 Receptor Antagonist |
| S100A12 | S100 calcium-binding protein -A12 |
| REG1A | Regenerating Family Member 1 Alpha |
| ECRG4 | Esophageal cancer-related gene-4 |
| CLDN4 | Claudin-4 |
| PAX5 | Paired Box-5 |
| LACTB1 | Lactamase beta like-1 |
| CORIN | Corin |
| SOST | Sclerostin |
| MFRP | Membrane Frizzled-Related protein |
| TMEM72 | Transmembrane-72 |
| SHOX2 | Short Stature Homeobox-2 |
| WFIKKN2 | WAP, Follistatin/Kazal, Immunoglobulin, Kunitz And Netrin Domain Containing 2 |
| SLC4A5 | Solute Carrier Family 4 Member 5 |
| LHX9 | LIM Homeobox Protein 9 |
| SLC13A1 | Solute Carrier Family 13 Member 1 |
| UCN3 | Urocortin-3 |
| TYMP | Thymidine Phosphorylase |
| FREM3 | FRAS1 Related Extracellular Matrix 3 |
| ESRRG | Estrogen Related Receptor Gamma |
| EYA4 | Eyes Absent Homolog 4 |
| SLITRK6 | SLIT and NTRK-like family member 6 |
| GUCY1A1 | Guanylate Cyclase 1 Soluble Subunit Alpha 1 |
| EDIL3 | EGF-like repeats and discoidin I-like domains 3 |
| CDKL2 | Cyclin Dependent Kinase Like 2 |
| PRKAR1A | Protein Kinase CAMP-Dependent Regulatory Type I Alpha |
| DDAH1 | Dimethylarginine Dimethylaminohydrolase 1 |
| SPON1 | Spondin-1 or F-spondin |
| RAD21 | RAD21 Cohesin Complex Component |
| MYO1B | Myosin 1B |
| RASGEF1 | RasGEF domain family, member 1A |
| FGF12 | Fibroblast Growth Factor 12 |
| ARSA | Arylsulfatase A |
| ATM | Ataxia-Telangiectasia Mutated |
| C1QA | A-chain polypeptide of complement subcomponent C1q |
| C3AR1 | Complement Component 3a Receptor 1 |
| C5AR1 | Complement Component 5a Receptor 1 |
| CASP1 | Caspase-1 |
| CCL2 | Chemoattractant Protein-1 |
| CD14 | Cluster of Differentiation 14 |
| CD300LF | CD300 molecule-like family member F |
| CD36 | Cluster of Differentiation 14 |
| CLCN3 | Chloride Voltage-Gated Channel 3 |
| CSF1 | Colony-Stimulating Factor 1 or Macrophage Colony-Stimulating Facto (M-CSF) |
| CSF1R | Colony-Stimulating Factor 1 Receptor |
| CYBB | Cytochrome b-245, beta chain. |
| DBI | Diazepam Binding Inhibitor, Acyl-CoA Binding Protein |
| DOCK10 | Dedicator of Cytokinesis 10 |
| F2RL1 | Coagulation Factor II Thrombin Receptor-Like 1 |
| GRN | Granulin Precursor |
| HTRA1 | High Temperature Requirement A Serine Peptidase 1 |
| IFNGR1 | Interferon Gamma Receptor 1 |
| ITGAL | Integrin Subunit Alpha L |
| ITGAV | Integrin Subunit Alpha V |
| ITGB1 | Integrin subunit beta 1 or CD29 |
| JAK2 | Janus kinase 2 |
| LGALS3 | Galectin-3 |
| LYN | LYN Proto-Oncogene, Src Family Tyrosine Kinase |
| MBP | Myelin Basic Protein |
| MERTK | MER proto-oncogene, tyrosine kinase |
| NR3C1 | Nuclear Receptor Subfamily 3, Group C, Member 1 |
| NR3C2 | Nuclear receptor subfamily 3 group C member 2 |
| NTF3 | Neurotrophin 3 |
| PDK4 | Pyruvate Dehydrogenase Kinase 4 |
| PLAT | Plasminogen Activator, Tissue Type |
| PLP1 | Proteolipid Protein 1 |
| PPARG | Peroxisome Proliferator-Activated Receptor Gamma |
| PTEN | Phosphatase and TENsin homolog |
| PTPRC | Protein Tyrosine Phosphatase, Receptor Type C |
| RGS10 | Regulator of G-protein Signaling 10 |
| RUNX1 | Runt-related transcription factor 1 |
| S100A9 | S100 calcium-binding protein A9 |
| SERPINE1 | Serpin Family E Member 1 |
| SERPIN1 | Makes alpha-1 antitrypsin |
| SPP1 | Secreted phosphoprotein 1 |
| SST | Somatostatin |
| STAT6 | Signal Transducer and Activator of Transcription 6 |
| TLR1 | Toll-like receptor 1 |
| TLR4 | Toll-like receptor 4 |
| TSPAN2 | Tetraspanin 2 |
| TYROBP | TYRO protein tyrosine kinase binding protein |
| CTSB | Cathepsin B |
| FTL | Ferritin Light Chain |
| RRM2 | Ribonucleotide Reductase Regulatory Subunit M2 |
| LPCAT1 | Lysophosphatidylcholine Acyltransferase 1 |
| CDK1 | Cyclin-dependent kinase 1 |
| NFE2L2/NRF2 | Nuclear Factor, Erythroid 2-Like 2, code NRF2 protein |
| SQSTM1 | Sequestosome 1 |
| NDUFS2 | Ubiquinone Oxidoreductase Core Subunit S2 |
| BACH1 | Transcription factor BTB and CNC homology 1 (Bach1) |
| FLVCR1 | Feline Leukemia Virus Subgroup C Receptor 1 |
| PRKC1 | Protein Kinase C Iota |
| TFRC | Transferrin receptor |
| SLC7A11 | Solute Carrier Family 7 Member 11 |
| ACSL4 | Acyl-CoA Synthetase Long-Chain Family Member 4 |
| ABCA4 | Retinal-specific ATP-binding cassette transporter |

